# Serine mistranslation induces the integrated stress response without accumulation of uncharged tRNAs

**DOI:** 10.1101/2024.02.04.578812

**Authors:** Hong Zhang, Jiqiang Ling

## Abstract

Aminoacyl-tRNA synthetases (aaRSs) are essential enzymes that support robust and accurate protein synthesis. A rapidly expanding number of studies show that mutations in aaRSs lead to multiple human diseases, including neurological disorders and cancer. Much remains unknown about how aaRS mutations impact human health. In particular, how aminoacylation errors affect stress responses and fitness in eukaryotic cells remains poorly understood. The integrated stress response (ISR) is an adaptive mechanism in response to multiple stresses. However, chronic activation of the ISR contributes to the development of multiple diseases (e.g., neuropathies). Here we show that Ser misincorporation into Ala and Thr codons, resulting from aaRS editing defects or mutations in tRNAs, constitutively active the ISR. Such activation does not appear to depend on the accumulation of uncharged tRNAs, implicating that Ser mistranslation may lead to ribosome stalling and collision.

## Introduction

Aminoacyl-tRNA synthetases (aaRSs) universally exist in all organisms and are essential enzymes playing critical roles in protein synthesis. AaRSs define the first step of protein synthesis by charging cognate tRNAs with amino acids to form aminoacyl-tRNAs (aa-tRNAs), which are delivered to the ribosome for nascent peptide formation (1-3). In some cases, mischarging of tRNAs by aaRSs occurs (4,5). Most organisms have evolved aaRS editing activities to deacylate the mischarged aa-tRNAs. The editing function is an essential checkpoint to ensure translation fidelity (6,7). An increasing number of studies show that mutations in aaRS genes result in various neurological diseases (e.g., Charcot–Marie–Tooth (CMT) disease and microcephaly) (8-11), developmental delay (12-15), and cancer (16-18). Other factors involved in tRNA biogenesis and tRNA modifications are also linked to multiple diseases (19-21). Revealing the physiological effects of aaRS mutations is thus critical to understanding the development of neurodegenerative diseases and searching for effective drugs to treat these diseases.

Mistranslation of the genetic code by deficient aaRSs is usually considered unfavorable for cell growth. Faithful translation of the genetic code into active protein is crucial for cell viability, as translation errors can lead to the accumulation of misfolded proteins and protein aggregates which are toxic to the cells (22,23). It has been shown that mutations in alanyl-tRNA synthetase (AlaRS) leading to an editing defect cause damage to neurons and cardiomyocytes (24). Mutations in the editing domain of yeast AlaRS and threonyl-tRNA synthetase (ThrRS) cause sensitivity to heat stress (25,26). However, translational infidelity may benefit bacteria under certain stress conditions (27-29), partially due to the activation of stress responses by moderate mistranslation.

Diverse stressful conditions activate the integrated stress response (ISR), which is an evolutionarily conserved signaling pathway that adapts cells to stresses (30). In mammalian cells, the ISR is mediated by four stress-sensing kinases (PERK, GCN2, PKR, and HRI) to reduce overall protein biosynthesis while allowing translation of specific genes to support adaptation to adverse environments (31). The ISR is also known as the general amino acid control (GAAC) signaling pathway in yeast (32-34). Amino acid starvation accumulates uncharged tRNAs, which bind to and activate Gcn2. Activated Gcn2 phosphorylates eukaryotic initiation factor 2 α-subunit (eIF2α) on Ser 51, which attenuates global translation and triggers the Gcn4-mediated amino acid starvation response (35). Recent work also suggests that ribosome collision may activate the ISR without notable accumulation of uncharged tRNAs (36,37). Interestingly, dominant CMT mutations in glycyl-tRNA synthetase activate the ISR by inducing ribosome stalling (38-40) and inhibiting Gcn2 and the ISR alleviates peripheral neuropathy in a mouse model (38).

Mutations in the editing sites of AlaRS and ThrRS are found to cause microcephaly and developmental disorders (13,15,41). Both AlaRS and ThrRS mischarge Ser onto tRNAs and require editing to remove the mischarged tRNAs. In our previous work, we show that an AlaRS editing-site mutation (C719A) leads to increased Ser mistranslation and activation of the ISR (25), but the underlying mechanism is unclear. In this work, we show that Ser misincorporation at Ala and Thr codons induces phosphorylation of eIF2α and *GCN4* expression. We further show that activation of the ISR by Ser mistranslation does not require an increase in uncharged tRNAs, implicating the involvement of ribosome stalling and collision in mistranslation-induced ISR.

## Results

### Transcriptome analysis of a ThrRS editing-defective yeast strain

We have recently shown that a ThrRS editing-defective yeast strain (*ths1-C268A*) is sensitive to heat stress (26). To understand the global transcriptome change caused by ThrRS editing defects, we performed RNA sequencing of the WT and *ths1-C268A* strains under heat stress. We grew the cells in YPD to log phase at 30 °C and shifted the cultures to 37 °C heat stress for 2 h before collecting cells for transcriptome analysis. Compared to the WT, multiple pathways were significantly changed in the editing defective ThrRS strain. Notably, the amino acid biosynthesis pathway was the most significantly upregulated (Fig. 1A). On the contrary, significantly downregulated pathways include rRNA modification, ribosome biogenesis, cell cycle, DNA replication and repair, TCA cycle and response to DNA damage (Fig. 1B).

**Figure 1.**
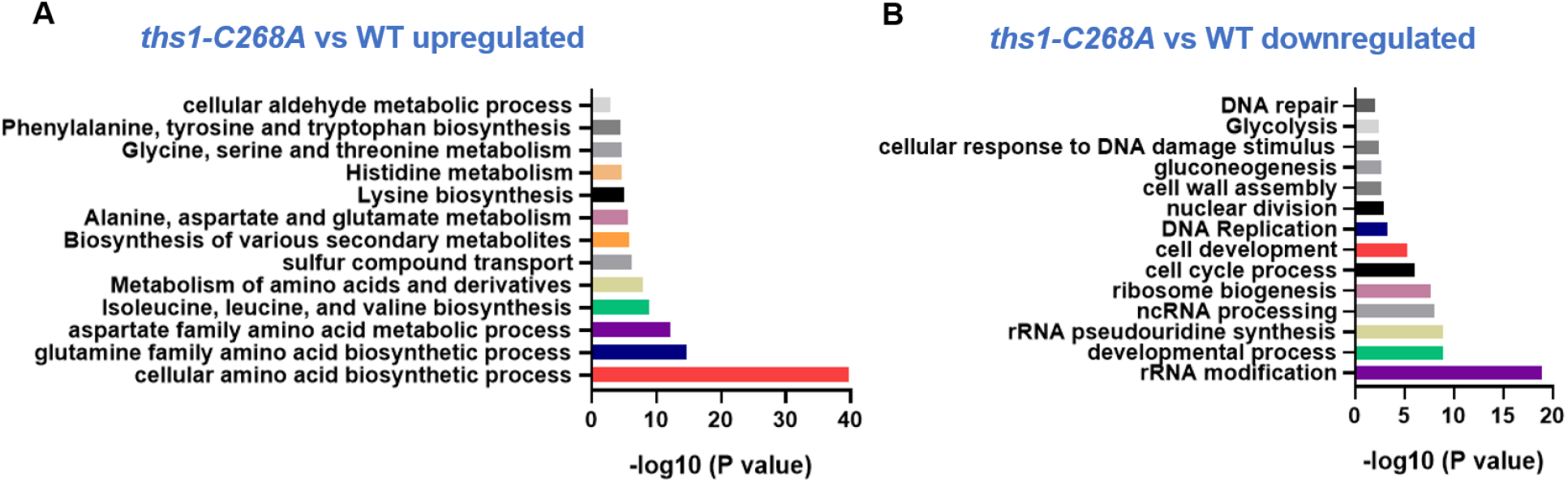
Transcriptome analysis of ThrRS editing-defective mutant *ths1-C268A* and WT under heat stress. Cells were grown in YPD to the log phase at 30 °C and shifted to 37°C for 2 h. Upregulated (A) and (B) downregulated pathways are shown. Three replicates were performed for each strain.

### ThrRS editing deficiency activates the ISR

In yeast, the amino acid biosynthesis pathway is activated by Gcn4 via phosphorylation of eIF2α (Fig. 2A). We transformed the *GCN4-LacZ* reporters pJD821 and pJD823 (positive control) into WT and *ths1-C268A* strains. Cells were cultured at 30 °C to the log phase, followed by 2 h of treatment at 37 °C before the β-galactosidase assay. As shown in Fig. 2B, *GCN4* expression was significantly increased by the *ths1-C268A* mutation, consistent with our RNA sequencing result. In yeast, Gcn2 is the only known kinase to phosphorylate eIF2α. We confirmed that activation of the ISR in *ths1-C268A* was dependent on Gcn2, as deleting the *GCN2* gene abolished *GCN4* expression in the *ths1-C268A* strain (Fig. 2C). As the Gcn4-mediated amino acid biosynthesis is induced in *ths1-C268A* under heat stress, we tested the role of Gcn4 on heat sensitivity. Deleting *GCN4* did not restore heat resistance in *ths1-C268A* (Fig. 2D), suggesting that other critical pathways are responsible for heat sensitivity.

**Figure 2.**
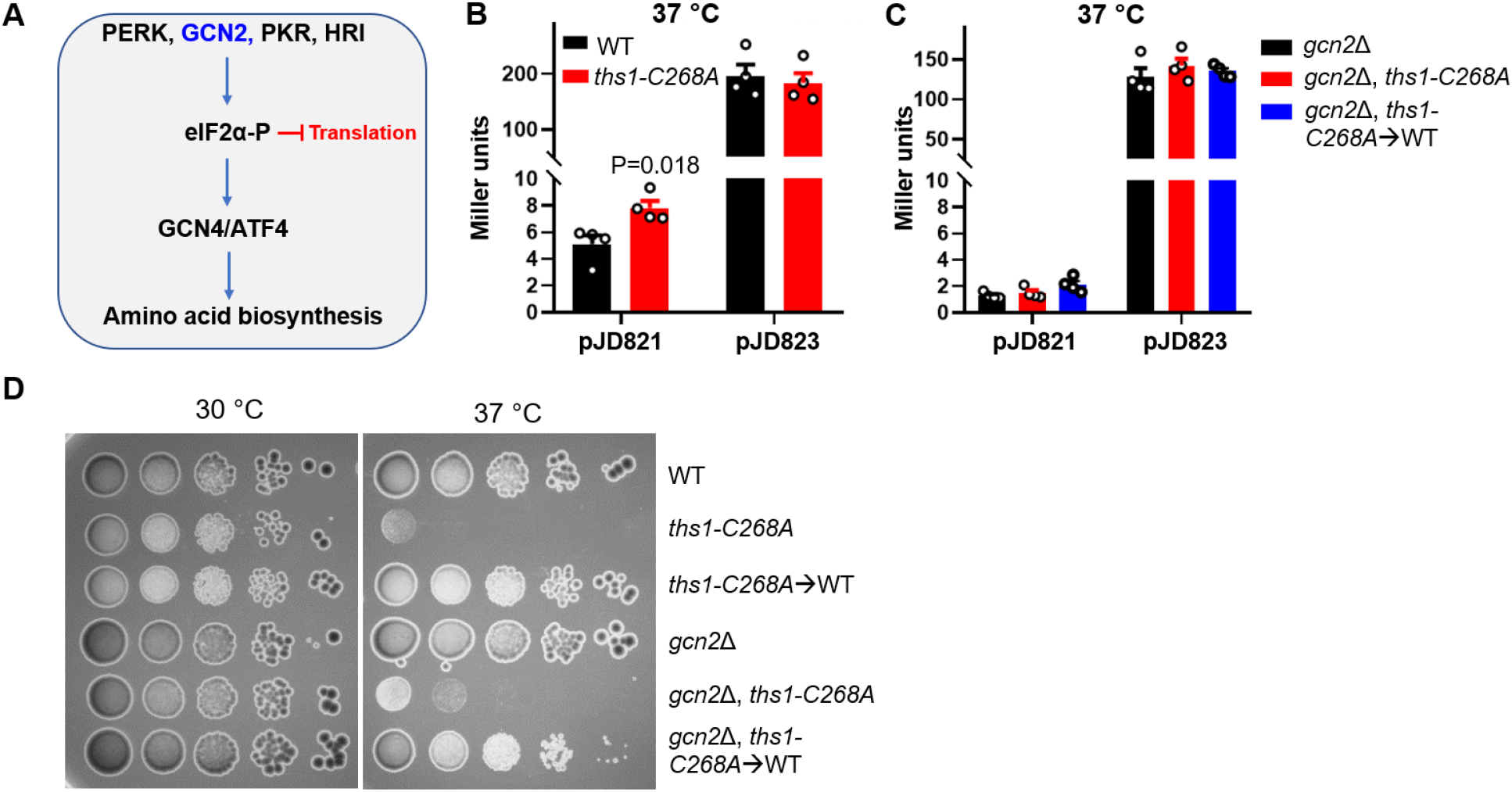
*ths1-C268A* mutation activates the integrated stress response under heat stress. (A) The ISR is activated by four protein kinases PERK, GCN2, PKR, and HR in mammals. GCN2 is the only kinase known to activate the ISR in yeast. The ISR attenuates global protein synthesis and activates amino acid biosynthesis via GCN4/ATF4. (B, C) Expression of *GCN4* shown by LacZ reporters. pJD821 contains all the regulatory elements for *GCN4* expression; pJD823 is a positive control without the regulatory elements. Each circle represents one biological replicate. Error bars represent one SD from the mean. The P values are determined using the unpaired t -test. (D) Yeast cells were grown to the log phase and spotted on YPD agar plates with 10-fold dilutions. The plates were incubated at 30 °C and 37 °C before imaging. Representative images of three biological replicates are shown.

We have previously shown that the *thr1-C268A* mutation decreases the stability of ThrRS at 37 °C in addition to abolishing the editing activity (26). A decreased level of aaRSs may lead to the accumulation of uncharged tRNAs and activation of the ISR. Because the stability of the ThrRS C268A mutant is not affected at the normal temperature for yeast (30 °C), we decided to test whether an editing defect alone is sufficient to induce the ISR at 30 °C. We found that the *thr1-C268A* mutation also activates the Gcn2-dependent ISR at 30 °C (Fig. 3). We further used the *thr1-C268A*⟶WT revertant to show that activation of the ISR is indeed due to the *thr1-C268A* mutation, but not off-target mutations in the CRISPR-engineered strain. Further addition of the His analog 3-aminotirazole (3-AT) increased *GCN4* expression in all three strains (Fig. 3C). Collectively, these results support that ThrRS editing deficiency, similar to AlaRS, activates the ISR in a Gcn2-dependent manner.

**Figure 3.**
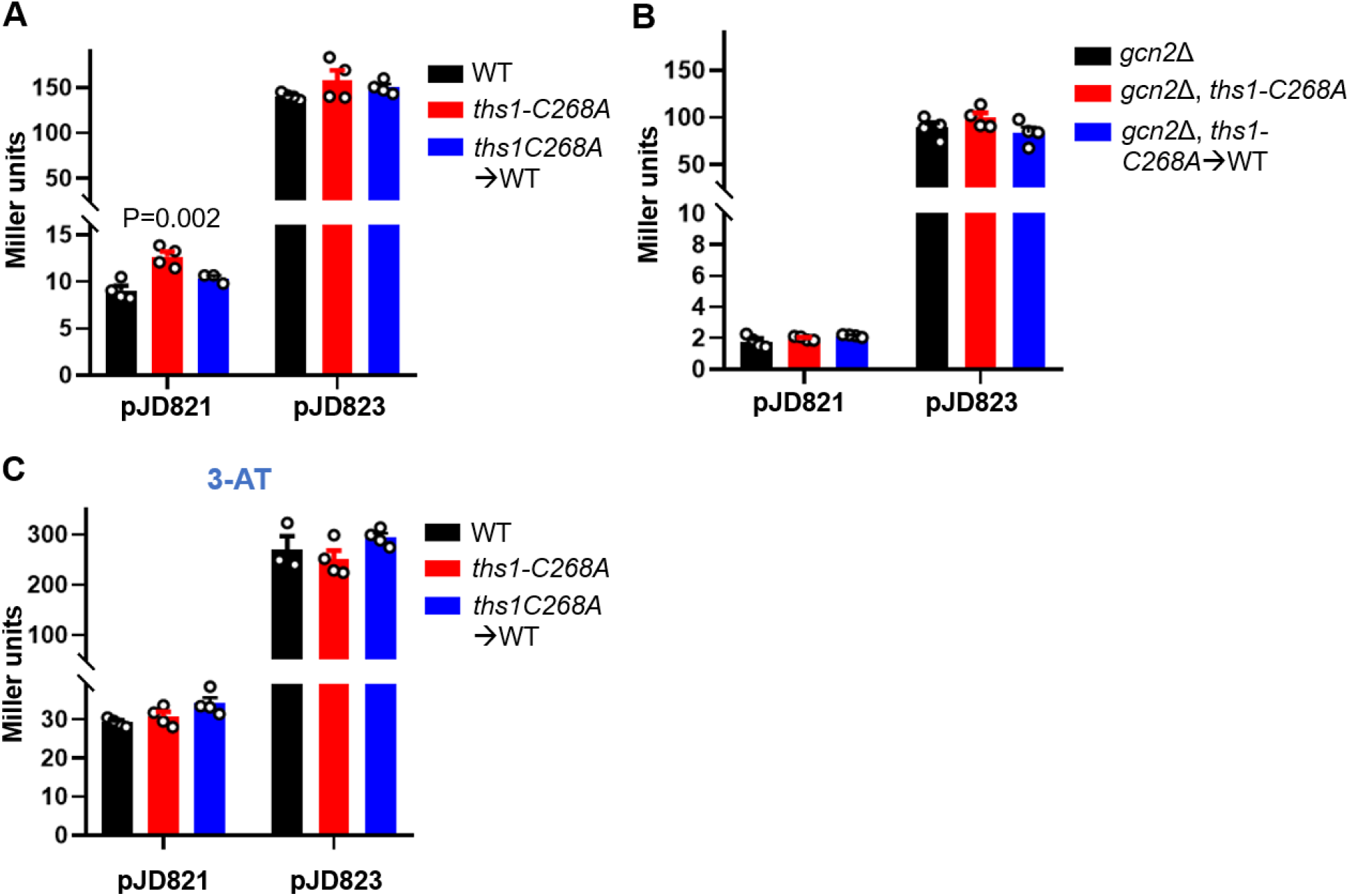
*ths1-C268A* mutation activates the ISR at normal temperature (30 °C). (A, B) Expression of *GCN4* shown by LacZ reporters. (C) Cells were treated with 100 mM 3-AT before the LacZ assay. Each circle represents one biological replicate. Error bars represent one SD from the mean. The P values are determined using one-way ANOVA.

### Severe Ser misincorporation at Thr codons by tRNA^Ser^ variants causes growth defects

In addition to aaRS editing defects, amino acid mistranslation can also result from tRNA variants, particularly in the anticodons. Anticodon variants of tRNAs are indeed frequently found in human populations (42), and mistranslating tRNAs are actively pursued as gene therapies for genetic diseases (43,44). To test whether mistranslating tRNAs affect fitness and activate the ISR, we mutated the anticodon of tRNA^Ser^ from AGA (WT) to AGT and TGT to recognize Thr codons instead of Ser codons, and expressed the tRNAs in WT yeast on a single-copy number plasmid (pRS315). Seryl-tRNA synthetase does not recognize the anticodon and attaches Ser to tRNA^Ser^ anticodon variants (45). Using a β-lactamase reporter assay (26), we found that expressing tRNA^Ser^_AGT_ increased the Ser misincorporation rate at the ACT Thr codon to ∼4% (Fig. 4A). Expressing tRNA^Ser^_AGT_ or tRNA^Ser^_TGT_ impaired growth at 30 °C and almost abolished growth at 37 °C (Figs. 4B-4E), suggesting that severe Ser mistranslation is harmful to cells even at normal temperatures.

**Figure 4.**
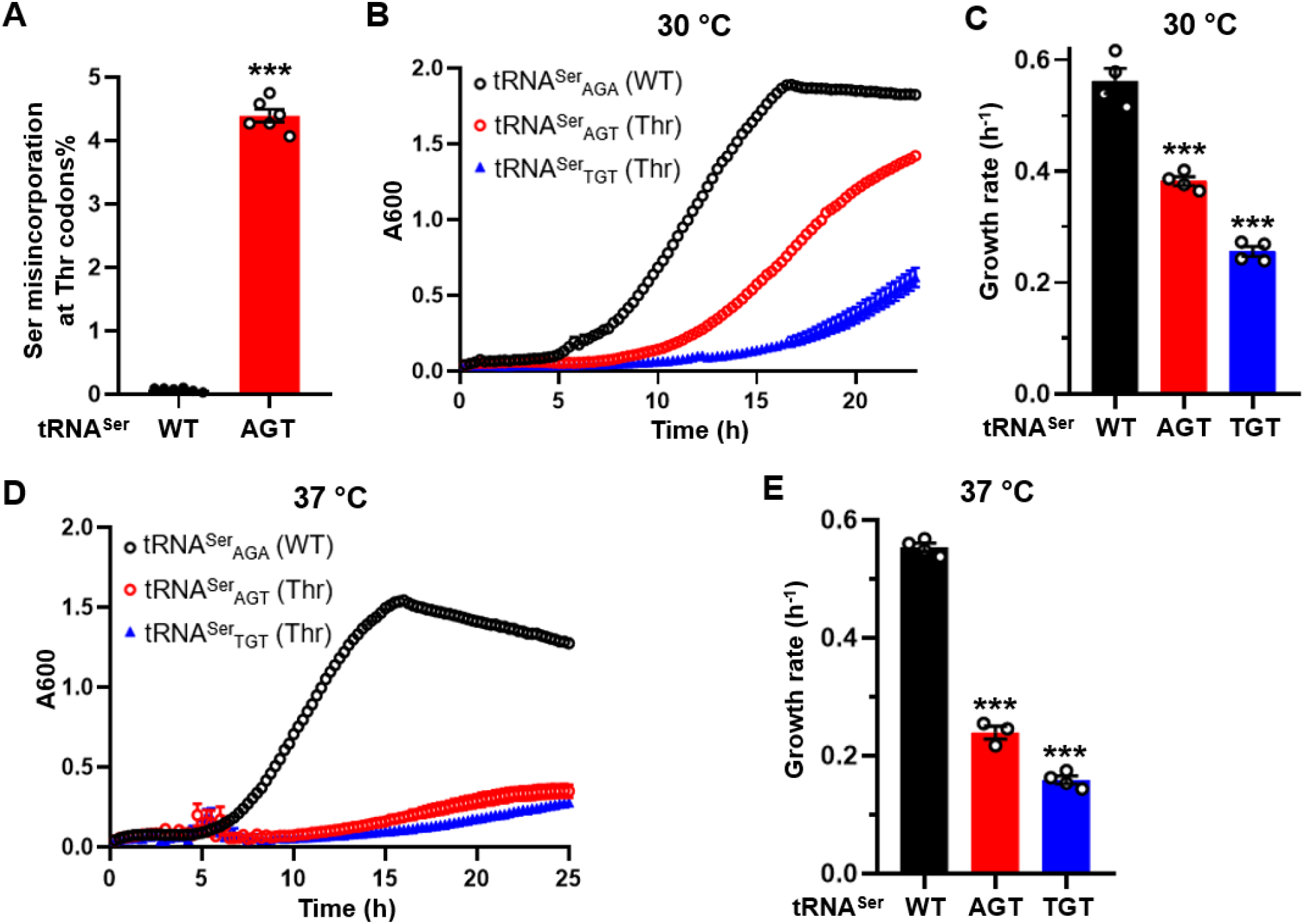
Serine mistranslation at Thr codons caused by expressing tRNA^Ser^ variants impair growth. (A) Ser misincorporation at Thr codon was determined using an S68T β-lactamase variant. Growth curves and rates of WT yeast carrying pRS315-tRNA^Ser^ variants at 30 °C (C, D) and 37 °C (E, F). (B, D) show the mean growth of at least three biological replicates with error bars indicating one SD. Each circle in (A, C, D) represents one biological replicate. B Error bars represent one SD from the mean. The P values are determined using one-way ANOVA. *** p < 0.001.

### Ser misincorporation activates the ISR without increasing uncharged tRNAs

We next tested whether tRNA^Ser^ variants activate the ISR. We selected tRNA^Ser^_AGA_ (WT), tRNA^Ser^_TGT_ (Thr), tRNA^Ser^_AGC_ (Ala), and the empty vector. Expressing the WT tRNA^Ser^_AGA_ did not affect growth compared with the vector control but expressing the mistranslating tRNAs decreased growth (Fig. 5A). Both tRNA^Ser^_TGT_ and tRNA^Ser^_AGC_ enhanced *GCN4* expression (Fig. 5B). Phosphorylation of eIF2α at Ser51 plays a central role in ISR signaling. As shown in Fig. 5C, the level of eIF2α-P in both mistranslating strains substantially increased. Deleting *GCN2* abolished the phosphorylation of eIF2α in all tested strains (Fig. 5C), confirming that mistranslation-activated ISR depends on Gcn2. Gcn2 is recruited to the ribosome by Gcn1 (37,46,47). Knocking out *GCN1* diminished eIF2α phosphorylation (Fig. 5D), suggesting that activation of the ISR upon Ser mistranslation requires Gcn2 to associate with the ribosome.

**Figure 5.**
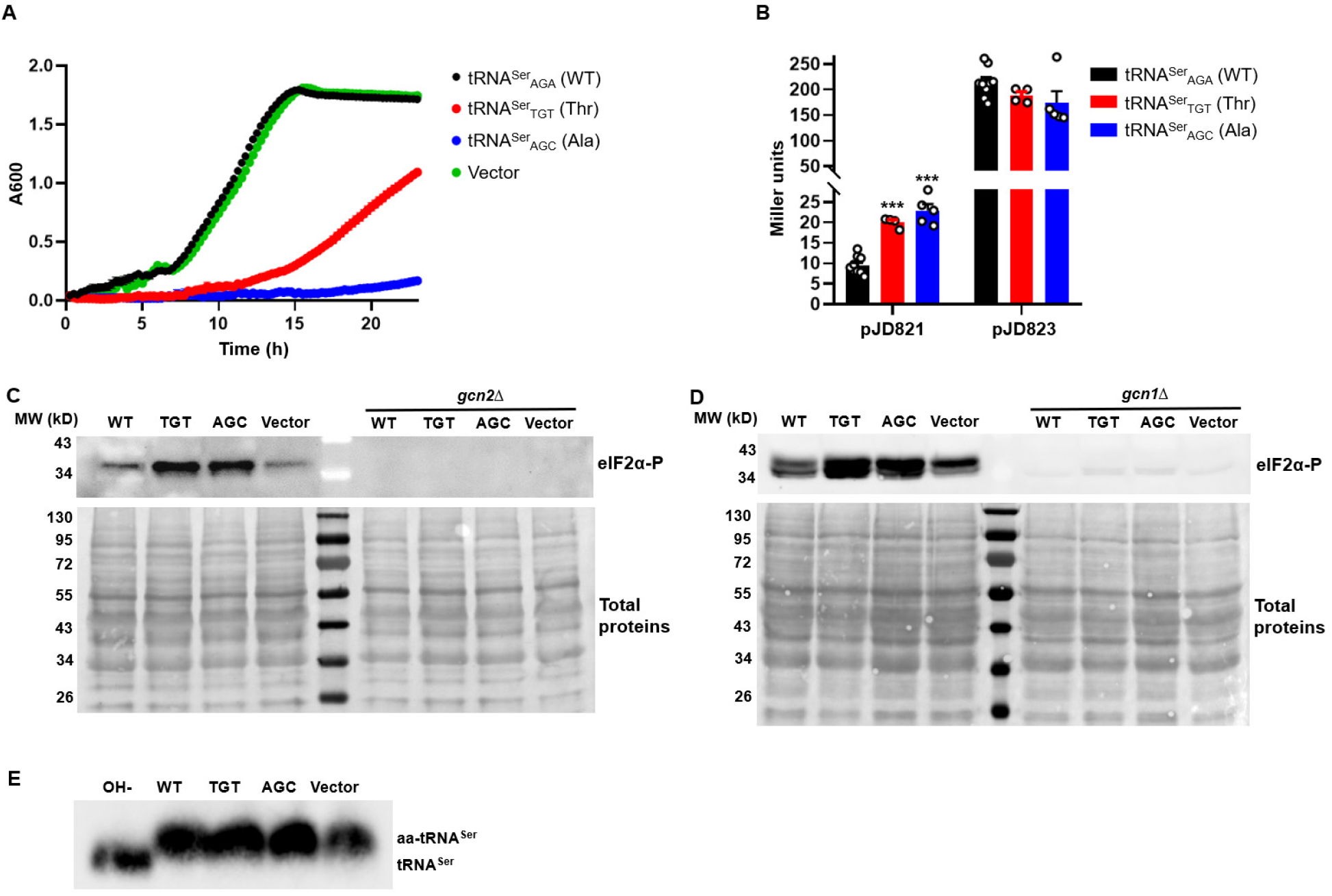
Mistranslating tRNA^Ser^ variants activates the ISR without increasing uncharged tRNA^Ser^. (A) Growth curve of WT yeast expressing tRNA^Ser^ variants in SD-Leu medium at 30 °C. The growth curves are the mean of at least three biological replicates with error bars indicating one SD. (B) Expression of *GCN4* shown by LacZ reporters in yeast expressing tRNA^Ser^ variants. Each circle represents one biological replicate. Error bars represent one SD from the mean. The P values are determined using one-way ANOVA. *** p < 0.001. (C, D) eIF2α phosphorylation detected by Western blot. Total proteins are revealed by Ponceau staining of the transferred membranes. (E) Acidic northern blot showing aminoacylated and deacylated tRNA^Ser^. OH-treatment deacylates the aa-tRNA. (C-E) show representative images of at least three biological replicates.

A possible mechanism of mistranslation-induced ISR is the accumulation of uncharged tRNAs. To test this, we used an acidic gel assay to probe the charging level of tRNA^Ser^. We found that tRNA^Ser^ was close to 100% aminoacylated in all four strains (Fig. 5E), indicating that Ser mistranslation promotes the ISR without elevating the level of uncharged tRNAs.

## Discussion

The central dogma explains the flow of genetic information from DNA through mRNA to proteins. Besides aminoacylation, around half of the aaRSs have evolved editing activities to ensure accurate protein synthesis (4,48). Given the central role of aaRSs in robust and faithful translation, it is not surprising that pathogenic mutations in aaRSs are increasingly associated with various human diseases, including peripheral neuropathies (8,9,49), brain development (50,51), neurodegenerative diseases like Alzheimer’s disease (AD) (52), autoimmune disorders (53), cancer (17,54), and cardiovascular diseases (55-58). Almost all cytosolic and mitochondrial aaRSs are implicated in the pathology of the human nervous system (51,59). CMT, a peripheral neurological disease characterized by muscle weakness and limb atrophy, was the first discovered neurological disorder to be linked to aaRS mutations (8). Cytosolic aaRS mutations leading to CMT are dominant and do not often cause aminoacylation defects (9). Recent studies using mice models and human cell lines suggest that chronic activation of the ISR plays a central role in CMT caused by aaRS mutations (38). CMT-causing glycyl-tRNA synthetase (GlyRS or GARS) mutants sequester tRNA^Gly^ and reduce the supply of Gly-tRNA^Gly^ to the ribosome, leading to ribosome stalling and activation of the ISR. Inhibiting ISR by deleting *GCN2* or using a small molecule inhibitor ISRIB remarkably alleviates the symptoms in mice (38). Overexpressing tRNA^Gly^ in GlyRS mutant flies and mice also suppresses the ISR and rescues the neuropathy phenotype (39). Whether ISR activation is responsible for peripheral neuropathies caused by other aaRS mutations remains to be determined.

In contrast to the dominant aaRS mutations affecting peripheral neurons, cytosolic aaRS mutations affecting the central nervous system are biallelic and recessive (9,50). Biallelic aaRS mutations identified in patients often decrease the aminoacylation efficiency or the stability of the mutant aaRSs (13,15,41,50). Decreased aminoacylation efficiency has been shown to activate the ISR in yeast and mammalian cells (25,60), likely through the accumulation of uncharged tRNAs. Multiple pathogenic mutations have also been mapped to the editing sites of AlaRS and ThrRS (13,15,41). Here we show that Ser misincorporation at Ala and Thr codons robustly activates the ISR. Interestingly, the tRNA charging level appears to be unchanged (Fig. 5E), raising the question as to how mistranslation activates the ISR. Recent studies reveal that ribosome stalling and collision can induce the ISR (36,37,61,62). Gcn2 recruitment to the ribosome depends on Gcn1 (46,47). Structural studies reveal that Gcn1 binds to stalled ribosomes (63). We show that deleting *GCN1* or *GCN2* abolishes phosphorylation of eIF2α in mistranslating strains (Fig. 5), leading us to speculate that Ser mistranslation results in aberrant translation elongation that leads to ribosome stalling. Whether and how this happens remain intriguing questions for future exploration.

## Experimental procedures

### Materials, media, and strains

All *Saccharomyces cerevisiae* strains used here were derivatives of BY4741. *Escherichia coli* DH5α grown in LB medium was used for molecular cloning. Gene knockout was generated by replacing the coding regions with the *His2* or *Leu* gene and verified by PCR. Yeast point mutation mutants and gene knockout strains were grown in the YPD medium (1% yeast extract, 2% peptone, 2% glucose). To induce Ser misincorporation into Thr or Ala codons, the anticodon of tRNA^Ser^ was mutated and cloned to pRS315. For strains harboring plasmids, they were grown in Synthetic Defined (SD) dropout medium (0.17% yeast nitrogen base, 0.5% ammonium sulfate, 2% glucose, and 0.14% amino acid dropout mix, -His, -Leu or -Ura).

### Transcriptome analysis

WT and *ths1-C268A* cells were grown at 30 °C to the mid-log phase and further cultured at 37 °C for 2 h. Total RNA was prepared using an RNA extraction kit (Qiagen). Library construction and Illumina sequencing were performed by Novogene.

### β-galactosidase assay

To test the expression of *GCN4*, yeast strains carrying pJD821 and pJD823 reporters (25) were grown in SD dropout media to the log phase at 30 °C. 700 µl of cultures were collected and resuspended in 700 µl Z-buffer (60 mM Na_2_HPO_4_, 40 mM NaH_2_PO_4_, 10 mM KCl, and 1 mM MgSO_4_), and cell density was measured by OD_600_. Cells were lysed by adding 100 µl Chloroform and 50 µl 0.1% SDS and vortexing for 15 s. The reaction was started by the addition of 0.2 ml prewarmed ONPG (o-nitrophenyl-β-galactoside, 4 mg/ml in Z-buffer), and terminated by adding 500 µl Na_2_CO_3_ (1 M). Cell debris was removed by centrifugation, and OD_420_ of the supernatant was determined using a platereader (Synergy HT, BioTek). The β-galactosidase activity was calculated according to the following equation: Miller units = 1000*OD_420_/(T*V*OD_600_).

### Spot assay

Yeast cells from single colonies were resuspended in YPD or SD-Leu, grown at 30 °C to saturation, and diluted 1:50 for continued growth to the log phase. Aliquots with serial dilutions (10^0^ to 10^−5^) were spotted on YPD or SD-Leu agar plates, which were incubated at 30 °C or 37 °C for 2 to 4 days before imaging.

### Growth and temperature sensitivity analysis

The yeast cells were grown in YPD or SD-Leu at 30 °C to saturation and 1:50 in YPD or SD-Leu in 96-well plate. Growth was performed in a microplate reader (Synergy H1, BioTek).

### β-lactamase assay

A β-lactamase assay was used to determine the Ser misincorporation rate in the yeast cells as described (25,26).

### Acidic northern blot

Total RNA of yeast cells was extracted using a hot phenol method. Pellets were dissolved in sodium acetate buffer (pH5.2). All samples were stored at -80 °C till use. Acidic northern blot was conducted as described (26).

### Western blot

Western blot was essentially performed as described (26). Yeast cultures were grown overnight, and total proteins were extracted using TCA precipitation. Samples were separated on 12% SDS-PAGE gels and transferred to nitrocellulose membranes, A standard Western blotting procedure was followed. The rabbit anti-eIF2α-P antibody (Thermo) with 1:1000 dilution, the mouse monoclonal anti-PGK1 primary antibody (Invitrogen) with 1:5000 dilution, and goat anti-mouse IgG-HRP secondary antibody (Invitrogen) with 1:5000 dilution were used. Nitrocellulose membranes were treated with ECL chemiluminescent substrate reagents (Bio-Rad) and visualized using a ChemiDoc Imaging System (Bio-Rad).

## Acknowledgments

This work was funded by the National Institute of General Medical Sciences (R35GM136213 to J.L.). We thank Drs. Jonathan Dinman (University of Maryland) and Thomas Dever (National Institute of Child Health and Human Development) for plasmids and helpful discussions.

## Author contributions

H.Z. and J.L. designed the project; H.Z. performed all the experiments and analyzed the data; H.Z. and J.L. wrote and proofread the manuscript.

## Notes

### Competing Interest Statement

The authors have declared no competing interest.

